# The impact of familiarity on cortical taste coding

**DOI:** 10.1101/2022.02.17.480922

**Authors:** Stephanie M. Staszko, John D. Boughter, Max L. Fletcher

**Affiliations:** Department of Anatomy & Neurobiology, University of Tennessee Health Science Center, Memphis TN USA 38163

## Abstract

The role of the gustatory region of the insular cortex in mediating associative taste learning, such as conditioned taste aversion, has been well studied. However, while associative learning plays a role in some taste behaviors, such as avoidance of toxins, taste stimuli are often encountered by animals in their natural environment without explicit consequences. This type of inconsequential experience with sensory stimuli has been studied in other sensory systems, generally with the finding that neuronal responses habituate with repeated sensory exposure. The present study sought to determine the effect of taste familiarity on population taste coding in mouse gustatory cortex (GC). Using microendoscope calcium imaging, we studied the taste responses of visually identifiable neurons over five days of taste experience, during which animals could freely choose to consume taste stimuli. We found that the number of active cells in insular cortex, as well as the number of cells characterized as taste-responsive, significantly decreased as animals became familiar with taste stimuli. Moreover, the magnitude of taste-evoked excited responses increased, and inhibited responses decreased with experience. By tracking individual neurons over time, we focused on taste coding in a subpopulation of “stable” neurons that were present on all days of the taste familiarity paradigm. The population-level response across these stable cells was distinct when taste stimuli were novel but became more intercorrelated among those taste stimuli mice willingly consumed as the stimuli became familiar. Overall, these results highlight the effects of familiarity on taste responses in gustatory cortex.

## Introduction

The insular cortex (IC) integrates environmental and interoceptive information, including taste, olfactory, somatosensory and viscerosensory input that all arrive from a number of sources (Allen et al., 1991; Hanamori et al., 1997, 1998; Shi and Cassell, 1998; Maffei et al., 2012; Gogolla, 2017; Blankenship et al., 2019; Gehrlach et al., 2020; Mizoguchi et al., 2020). Taste-responsive neurons in the IC, comprising “gustatory cortex” (GC), appear to be largely unnecessary for the expression of basic taste recognition behaviors (i.e., taste reactivity or discrimination) but instead are implicated in the organization of feeding-and taste-related learned behaviors (Braun et al., 1982; Oliveira-Maia et al., 2012; Lin et al., 2015; Reilly, 2018; Schier and Spector, 2019; Bouaichi and Vincis, 2020; Staszko et al., 2020; Boughter and Fletcher, 2021; Stern et al., 2021). In fact, studies have shown that the activity of taste-responsive neurons in GC is modulated following different types of learning, including cue-taste association (Samuelsen et al., 2012; Mazzucato et al., 2019) and conditioned taste aversion (CTA) (Yasoshima and Yamamoto, 1998; Accolla and Carleton, 2008; Grossman et al., 2008; Lavi et al., 2018). These learned events impart salience on taste stimuli; for example, the association of taste with gastric malaise in CTA produces subsequent aversion, i.e., taste now signals the animal that they are consuming a potentially toxic food. However, not all ingestive behaviors are based on, or affected by, associative learning. It is likely that most gustatory encounters occur in the absence of any overt ingestive consequences. For example, animals become familiar with novel tastes over successive encounters. This process of familiarization is important to the animal both in driving future ingestive decisions (Domjan, 1976; Harder et al., 1989; Mura et al., 2018) as well as impacting subsequent learning (Lubow, 2009; Flores et al., 2016; Flores et al., 2018; Flores et al., 2022).

Relatively few studies have explored the neural correlates of taste familiarization, and those have reported somewhat conflicting results. Expression of neuronal activity markers such as the immediate early gene c-Fos, or pERK (phosphorylated extracellular regulated kinase), is increased in GC and other taste-related forebrain areas following exposure to novel taste stimuli and subsequently decreases with repeated experience (Koh et al., 2003; Lin et al., 2012; Bamji-Stocke et al., 2018; Kayyal et al., 2021). In contrast, a study of multiunit activity in the GC in rats actively sampling a taste stimulus over several days found that the average response to the stimulus actually increased as the taste became familiar, although this change was only apparent in the late phase of the response (Bahar et al., 2004). More recently, Flores et al. (Flores et al., 2022) conducted recordings in awake rats and described both an increase in taste discriminability and enhancement of the late-phase palatability response in GC neurons, as animals were exposed to a pair of tastants and water over 3 days.

Here, we combined in-vivo calcium imaging with miniaturized microendoscopes (miniscopes) to image GC neurons over multiple days of taste experience to investigate changes in GC activity as taste stimuli shift from novel to familiar. In a change from prior studies, which used 1-3 stimuli, we gave mice access to a larger panel of water and 6 novel stimuli representing each basic taste quality over a 5-day period. Moreover, this imaging approach allows for the quantification of large numbers of active neurons across time, as well as the ability to assess response magnitude in individual neurons. Overall, we found a decrease in both the number of active and taste-responsive neurons in GC across five days of taste exposure. Further, we find that mean taste-evoked excitatory responses increase and suppressed responses decrease. To further explore the role of familiarization on GC taste coding, we compared population response correlations for each taste stimulus on each day and found that GC taste representations for actively-licked compounds became more similar with familiarization.

## Methods

### Animals

Adult male (n=3) and female (n=6) C57BL/6J mice (The Jackson Laboratory) were used for all miniscope experiments. Age-matched mice (n=8) were used as non-surgical controls. Animals were housed on a 12-hour light/dark cycle and were group-housed until lens implant surgery, at which point they were individually housed to protect lens integrity. Non-surgical controls were similarly singly housed throughout behavioral experiments. All procedures were approved by the University of Tennessee Health Science Center Institutional Care and Use Committee.

### Virus Injections

Adult mice (4-6 weeks of age) were anesthetized using isoflurane (4-5% induction, 1-2% maintenance) and positioned in a stereotaxic apparatus (David Kopf Instruments). Carprofen (5.0 mg/kg, subcutaneous, Henry Schein) was administered prior to any surgical procedures. Following preparation of the surgical area, an incision was made from Bregma to lambda, and a craniotomy was formed above gustatory cortex (anterior +1.1 mm, lateral 3.3mm, and ventral 1.75 mm relative to Bregma) using a dental drill (Osada Inc). 500 nL of AAV1.Syn.GCaMP6s.WPRE.SV40 (Addgene) was injected at a speed of 15 ul/sec using a Nanoject II (Drummond Scientific). The micropipette was removed 10 minutes after virus injection, at which point the skin was closed with suture. Animals were allowed to recover for two weeks prior to the gradient refractive index (GRIN) lens implant.

### GRIN Lens Implantation

Animals were anesthetized with isoflurane and held in the stereotaxic apparatus using non-rupturing ear bars. Carprofen (5.0mg/kg) and dexamethasone (0.2mg/kg) were administered subcutaneously prior to surgical procedures. Following preparation of the surgical area, the skin was removed from the surface of the skull. A 4mm long GRIN lens (either 0.5mm or 1mm diameter) (Inscopix) was stereotaxically implanted into insular cortex (anterior +1.3mm, lateral 3.3mm, ventral 1.75 mm relative to Bregma) using a custom holder. The lens was first secured using cyanoacrylate glue, and then the entire skull was covered with dental cement (Ortho-Jet, Lang Dental) to seal the surgical area and create a head cap. The surface of the lens was covered with kwik-sil (World Precision Instruments) to prevent damage prior to baseplate surgery. Animals were given antibiotic food (Uniprim, Fisher Scientific) and a daily subcutaneous injection of carprofen/dexamethasone for five days following surgery. Animals were then given approximately 6-8 weeks of recovery to allow for optimal healing beneath the imaging window. To attach baseplates, animals were anesthetized with isoflurane and placed back in the stereotaxic apparatus. A 3D printed holder (ONE Core) mounted to the stereotaxic allowed for the placement of the baseplate. The baseplate was attached to the skull using dental cement. A custom lightweight metal head bar was also attached to the head cap using cyanoacrylate glue. The head bar allowed for brief fixation to ensure repeatable placement of the miniscope over multiple days of imaging.

### Davis Rig Acclimation and Training

All behavioral procedures were conducted in water-restricted mice using a contact lickometer (Davis MS-160, DiLog Instruments). The lickometer consists of a small chamber with 3 Plexiglas walls and a fourth wall made of stainless steel, the latter of which contains a stimulus port through which a mouse can access taste solutions. Access to the port is controlled by a computer-operated shutter. Stimulus bottles are held on a motorized tray and can be driven into position opposite the stimulus port for an ensuing trial. In these experiments, stimuli were presented at room temperature in 20 ml glass bottles with stainless-steel sipper tubes. Mice were trained and tested in the lickometer over a period of 2-3 weeks. Prior to being placed in the lickometer each day, animals were briefly head-fixed to allow for consistent placement of the miniscope field of view. On day 1, mice were placed in the lickometer with the shutter closed for 20 minutes to acclimate to the test chamber. After this session, access to water in the home cage was removed, beginning the water restriction schedule. Throughout both training and testing, mice received most of their daily fluid intake in the lickometer; body weights of water-restricted animals were monitored daily, and supplemental water was given as necessary to maintain 85% of the animal’s original body weight. On day 2, mice received 20 minutes of free access to water in the lickometer through the stimulus port (sipper tube training). On days 3-5, mice underwent trial training in the lickometer, in which they initiated up to 28 brief trial presentations of water (5 s duration), with an intertrial interval (ITI) of 7.5 seconds. The final training day involved increasing the ITI to 60 s to allow for a distinct separation of trials during subsequent calcium imaging analysis.

### Tgste Fgmiligrizgtion

Following lick training, animals were tested for 5 days with a multi-taste panel. One mouse was removed due to a Davis Rig malfunction on day 3 of testing, leaving the total number used for analysis at n=9. A group of control mice who did not undergo GRIN lens implantation (n=8) were tested without miniscopes to ensure the scopes did not alter taste behavior. Taste stimuli included 0.5 M sucrose, 0.2% w/v sodium saccharin, 0.3 M NaCl, 0.02 M citric acid, 0.01 M quinine hydrochloride, and 0.01 M monopotassium glutamate mixed with 0.0001 M inositol monotriphosphate. Stock solutions were freshly prepared at the start of each day using reagent-grade chemicals mixed in filtered water. Concentrations were chosen based on our previous imaging study (Fletcher et al., 2017). For each test session (1 per day), the multi-taste panel consisted of 5s trials of each stimulus, interspersed with 5-second water “rinse” trials, with an ITI of 60 seconds. The stimuli were presented randomly within 2 blocks of trials, for a total of 12 stimulus and 12 water trials. Both the number and timing of licks to each stimulus were recorded in the lickometer software. For all analysis, we only considered the first block of trials, as licks in the second block tend to diminish with attenuation of thirst. Licking was quantified as the total number of licks during each trial.

### Miniscope Imaging

Imaging was conducted using V3 miniscopes (www.miniscope.org) (Cai et al., 2016) A custom-built commutator (ONE Core) prevented tangling of the miniscope’s coax cable during behavior. All videos were recorded at 15FPS using miniscope software. Corresponding behavioral data were collected using a webcam (Logitech) mounted above the Davis Rig. This video was used to manually align calcium imaging and behavioral data in Fiji (version 1.53f). Gain, exposure, and LED intensity varied between animals but were held constant across imaging days. Imaging sessions were spatially down-sampled 2x in Fiji to increase computational efficiency. All subsequent pre-processing was conducted using the Python (3.6) implementation of CaImAn (Giovannucci et al., 2019). Files were then motion-corrected using rigid motion correction. Minimum correlation and peak-noise-ratio values for constrained non-negative matrix factorization (CNMF) were determined by mouse based on summary images. After running the CNMF algorithm, CNMF outputs were further filtered by applying a minimum signal-to-noise ratio of 2.0, meaning the fluorescence signal had to reach a value of at least 2x the determined “noise” value for that cell to be accepted. Accepted and rejected components were evaluated manually, and CNMF threshold values were adjusted, if necessary until false positives and negatives were minimized based on data visualization. Accepted components are categorized as active cells in subsequent analyses. Deconvolved traces were used for all downstream calcium imaging analyses. Cell spatial footprints were extracted and aligned across imaging sessions using the MATLAB (R2017b, Mathworks) implementation of CellReg (Sheintuch et al., 2017). Multiday cell registration was completed based upon recommendations of CellReg probabilistic modeling. Alignment matrixes were manually evaluated using ROI visualization, and maximal distance shifts were adjusted, if necessary to optimize cell tracking.

Further analysis of deconvolved traces was conducted in R studio (Version 1.2.5003) using custom R scripts. Calcium traces were first aligned based on CellReg assignments. Traces were then temporally cropped to include 5 seconds prior to the licking through 20 seconds post licking to reduce data size. Taste-evoked change in fluorescence (ΔF) values were then calculated by subtracting the mean fluorescence evoked 2 seconds prior to licking from the mean fluorescence evoked during a two-second window surrounding the peak fluorescence during the licking period (Fletcher et al., 2017). As fluorescence data extracted from CaImAn has already been scaled by a background fluorescence value, ΔF, rather than ΔF/F, was calculated to describe neuronal responses (Giovannucci et al., 2019). Responsive cells were categorized as having at least a +/-2.5 SD change in fluorescence (ΔF) compared to the 2 second pre-licking baseline period and having a ΔF value greater than one.

### Analyses

Analysis was conducted on 1889 neurons recorded across all sessions and mice. For hierarchical cluster analysis and correlation analysis, taste responses for each cell were normalized to the maximum taste-evoked response for that cell. Responsive cells from all mice were pooled by day. Entropy (H), a common measure of breadth of tuning used in taste recordings, was used to evaluate changes in cell tuning across days (Smith and Travers, 1979).

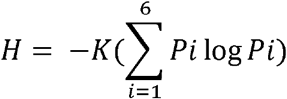

Pi represents the proportional excited response to each of the 6 tastes per cell, and K is a scaling constant (K = 1.283 for 6 tastants). Entropy value (H) ranges between 0 and 1, where 0 corresponds to a neuron that responds exclusively to a single stimulus (narrowly tuned) and 1 represents a neuron that responds equally to all stimuli (broadly tuned).

To better compare breadth of tuning to studies in other sensory systems, we also calculated each cell’s lifetime sparseness (S_L_) (Rolls and Tovee, 1995; Willmore and Tolhurst, 2001; Poo and Isaacson, 2009).

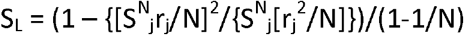

Where rj is the response of a neuron to tastant j and N is the total number of tastants. S_L_ ranges between 0 and 1, where 1 corresponds to a neuron that responds exclusively to a single stimulus (narrowly tuned) and 0 represents a neuron that responds equally to all stimuli (broadly tuned).

Statistical analyses for behavioral and imaging data were conducted using GraphPad Prism. Two-way repeated-measures ANOVA was used to test for differences in lick ratio between non-surgical control (n=8) and miniscope (n=8) animals across five taste days for each taste quality. Counts of active or taste-responsive cells were normalized to the first taste day to account for variations in the number of cells between animals. Changes in number of active cells across days were analyzed using the nonparametric repeated measures Friedman’s test along with Dunn’s multiple comparisons post hoc tests. Differences in mean taste evoked responses were analyzed using one-way ANOVA and unpaired t-tests. All graphs plot mean +/− SEM.

### GRIN Lens Placement Verification

At the conclusion of imaging experiments, animals were anesthetized with ketamine/xylazine (100/10 mg/kg IP) and transcardially perfused using 4% paraformaldehyde (PFA). Following perfusion, the brain was further postfixed in PFA for seven days to improve fixation and delineation of the imaging window. Brains were then removed, cryoprotected, and sectioned in 40 um thick serial sections using a freezing microtome. Sections were mounted on slides and imaged using a Nikon Eclipse 90i fluorescent microscope (Nikon Instruments Inc., Melville, NY USA) equipped with a digital camera and imaging software. GRIN lens placements were plotted on schematic section diagrams (Supp Fig. 1).

## Results

We used a combination of miniaturized microscopes and viral expression of the calcium indicator GCaMP6s to study how taste experience alters neuronal activity in the gustatory cortex of awake, freely behaving mice. To address this question, we allowed animals to freely lick taste stimuli and recorded calcium transients over the duration of five taste experience days. Figure 1 shows a typical GRIN lens placement within GC **(Figure 1A),** an example recording **(Figure 1B),** and identified soma **(Figure 1C).** The location of the GRIN lens for each mouse used in the study was verified to be in GC **(Supp. Figure 1)**. As miniscope recordings of GC have not been previously performed, we first characterized basic taste responses using day 1 recordings from all mice (n=9, 2350 cells). Cells within GC displayed robust taste responses to the tastants of our panel. **Figure 1D** shows example recordings of calcium traces taken from the same mouse during taste sampling.

**Figure 1.**
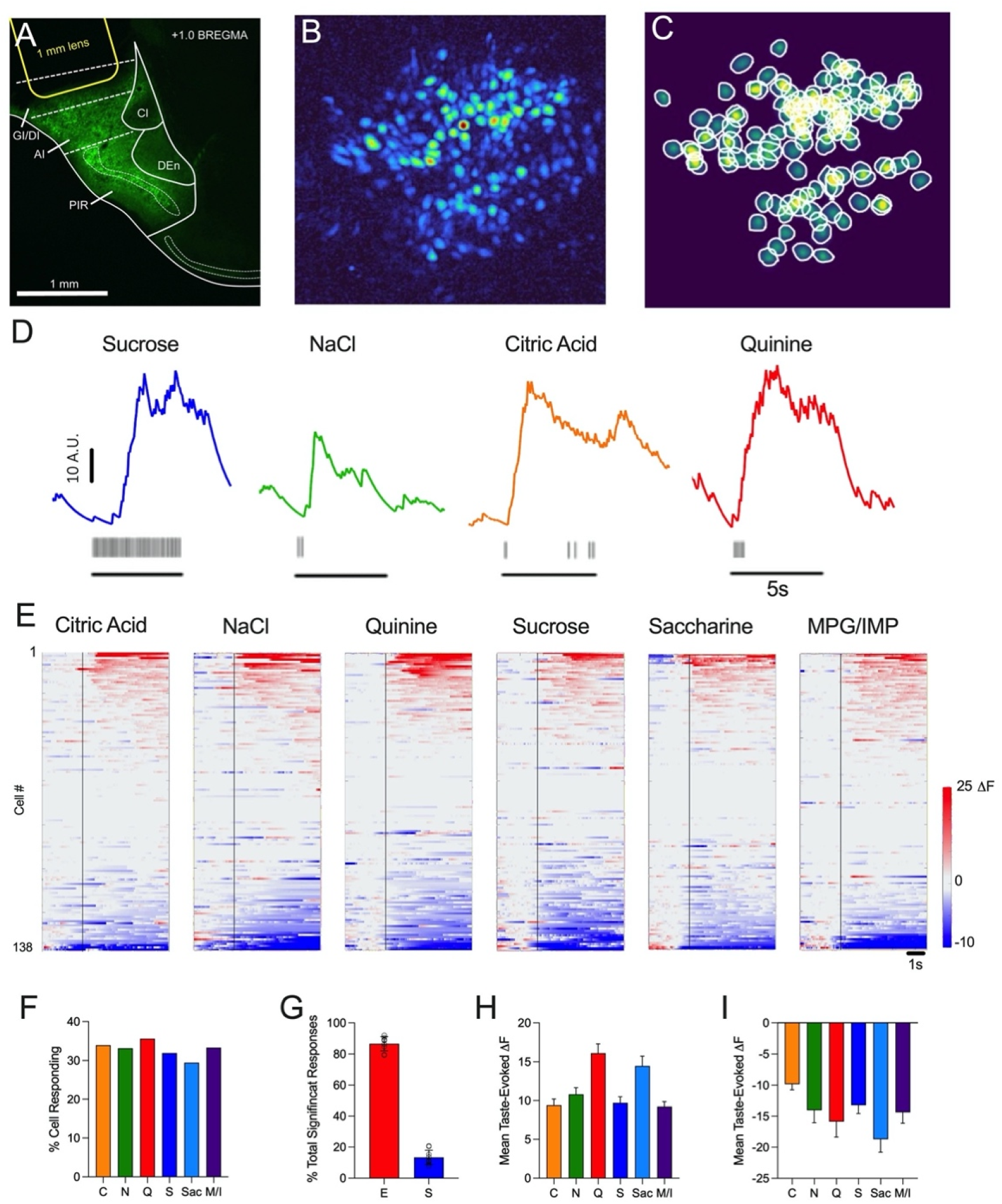
A. Coronal section through mouse brain showing an example of post imaging location verification, including GCaMP6 neuronal labeling. The space corresponding to the GRIN lens implantation is denoted by a yellow outline. B. C. D. Example traces from different GC neurons in response to licking of taste stimuli (tick marks under trace; line denotes the time when the mouse had access to spout). E. Normalized taste-evoked responses from all cells from an individual mouse before and during (vertical line) licking of 6 taste stimuli. Each row in the heat map represents a cell; red responses are excited, blue suppressed. F. Percentage of cells responding to each taste stimulus on day 1. G. Mean percentage of excited and suppressed cells in all mice on day 1 (n=9). H. Mean fluorescence change (excited) evoked by each taste across the cell population. I. Mean fluorescence change (suppressed) evoked by each taste across the cell population. GI/DI = granular, dysgranular insular cortex; AI = agranular insular cortex; PIR = piriform cortex. Error bars = SEM.

On average, we recorded from 261 ± 43 cells per mouse. Example recordings before and during taste presentations from a single animal (n=138 cells) are presented in **Figure 1E.** Taste-evoked excited responses in these heat maps are shown in red, and taste-evoked suppressed responses are shown in blue. Across all mice, we found 83% of recorded cells (n=1184) to be taste responsive as defined by having a significant response (excited or suppressed) to at least one of the six tastants. When pooled across all mice, we found an equal representation of responses as the percentage of cells responding to each was similar (Citric acid: 34%, NaCl: 33.2%, Quinine: 35.6%, Saccharine: 29.4%, Sucrose: 31.9%, MPG/IMP: 33.3%) **(Figure 1F).** Similar response representation was seen across individual mice, with each taste evoking similar percentages of cells. We did not observe overrepresentation of any single taste stimulus in any of the mice.

Within the total population of cells, 86.6% of the responses were excited, and 13.4% were suppressed **(Figure 1G).** Additionally, we found the mean taste-evoked DF responses to be similar across tastes for both excited and suppressed populations **(Figure 1H, I).** To quantify breadth of tuning, we used two different common methods, entropy and lifetime sparseness. Entropy values range from 0 and 1, where 0 corresponds to a narrowly tuned neuron, and 1 represents a broadly tuned neuron. In our data set, we found the mean entropy was 0.27 ± 0.01. To compare our findings to studies of cell tuning in other sensory systems, we also calculated the life sparseness for each neuron. This value also ranges from 0 and 1, but in this case, a value of 1 indicates a neuron that responds to a single stimulus, and a value of 0 indicates a neuron that responds to all stimuli. The mean sparseness was 0.86 ± 0.01.

With the exception of quinine, mice readily consumed all tastes each day **(Figure 2).** For each tastant, we performed a repeated-measures ANOVA across days to determine if licking changed with experience. For most tastes, we found no differences between lick numbers across days, suggesting that experience did not alter taste preferences. We did find a significant effect of day for both saccharine (F(1.80, 14.4)=4.70, p=0.0298) and NaCl (F(1.64, 13.2)=4.70, p=0.0019). Of the two tastants, post hoc Dunnett’s multiple comparisons tests revealed significant differences in NaCl licking between days 1 and 3, days 1 and 4, and days 1 and 5 (p<0.05), and no differences between day 1 and subsequent days of saccharine licking. Importantly, we demonstrate that the surgical procedures and miniscope-related handling do not affect normal consumption of basic taste stimuli, as miniscope-wearing mice were very similar to non-surgical controls in their lick ratios over five days of taste experience **(Supp. Figure 2).** Post-imaging reconstruction (Figure 1, Supp Figure 1) indicated that GRIN lenses were centered in either the granular (GI) or dysgranular division (DI) of insular cortex (or both) and were placed approximately within an anterior-posterior range from +1.2 – +0.5 mm relative to Bregma. This collective location corresponds with the gustatory representation described in previous studies with mice, although taste-responsive cells are also found in insular cortex more ventrally, anteriorly, and posteriorly (Chen et al., 2011; Fletcher et al., 2017; Levitan et al., 2019; Chen et al., 2021). The 1 mm GRIN lenses also typically extend across cortical layers (i.e., lateral to medial); however, it is not possible to determine for certain which layers are imaged in each mouse as this depends not only on the positioning of the lens but also on the extent of GCaMP expression in the region.

**Figure 2.**
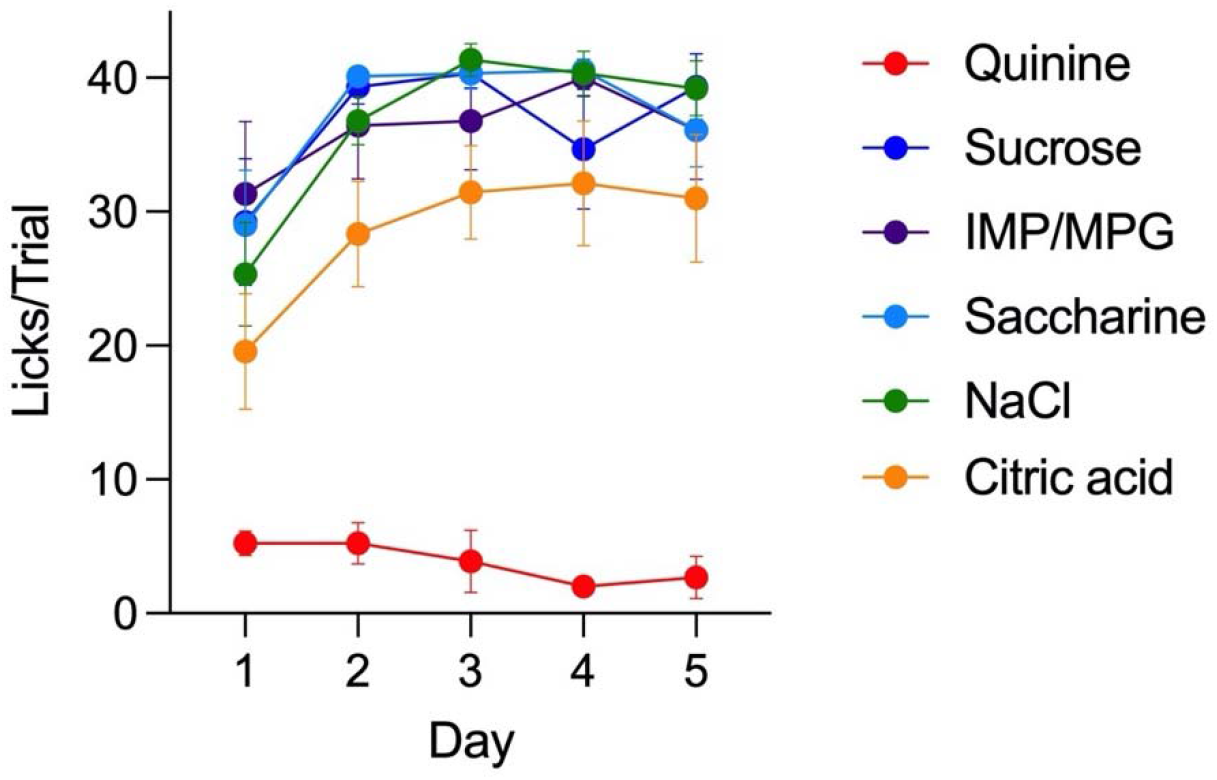
Mean licks per trial for each taste across 5 days of behavior. Error bars = SEM.

To investigate changes in GC activity related to familiarization, we first compared the number of total cells recorded each day from each mouse across the 5 days of taste presentations. Example traces taken from trials of each taste in one mouse on day 1 and day 5 are shown in **Figure 3A;** it is apparent that there were fewer active cells on day 5. Overall, we found a significant decrease in the number of active cells across days (D1: 100%, D2: 83.71 ±6.25%, D3: 88.58 ± 5.72%, D4: 86.20 ± 7.96%, D5: 77.36 ± 5.22%) (Friedman’s test, χ2(5) = 10.04, p = 0.0398) (n=8 mice). Post-hoc analysis with Dunn’s multiple comparison test revealed a significant difference between days 1 and 5 (p=0.0082). **(Figure 3B).** We also observed a similar drop in the number of taste responsive cells across days (D1: 100%, D3: 89.61 ± 7.42%, D5: 77.23 ± 4.04%) (Friedman’s test, χ2(3) = 10.89, p = 0.0029). Post-hoc analysis with Dunn’s multiple comparison test revealed a significant difference between days 1 and 5 **(p=0.0019) (Figure 3C).** The drop in active cells across days was not simply due to a reduction in the number of taste-responsive cells, as the ratio of the latter to the former did not significantly differ across days (Friedman’s test, χ2(3) = 0.22, p = 0.9712) **(Figure 3D).** To determine if the drop in cells was related to the taste experience, we compared the number of active cells observed across a similar period of water training days in a subset of mice (n=5). We found no statistical differences in the number of active cells across water days (Friedman’s test, χ2(3) = 1.60, p = 0.5216) **(Figure 3E)**.

**Figure 3.**
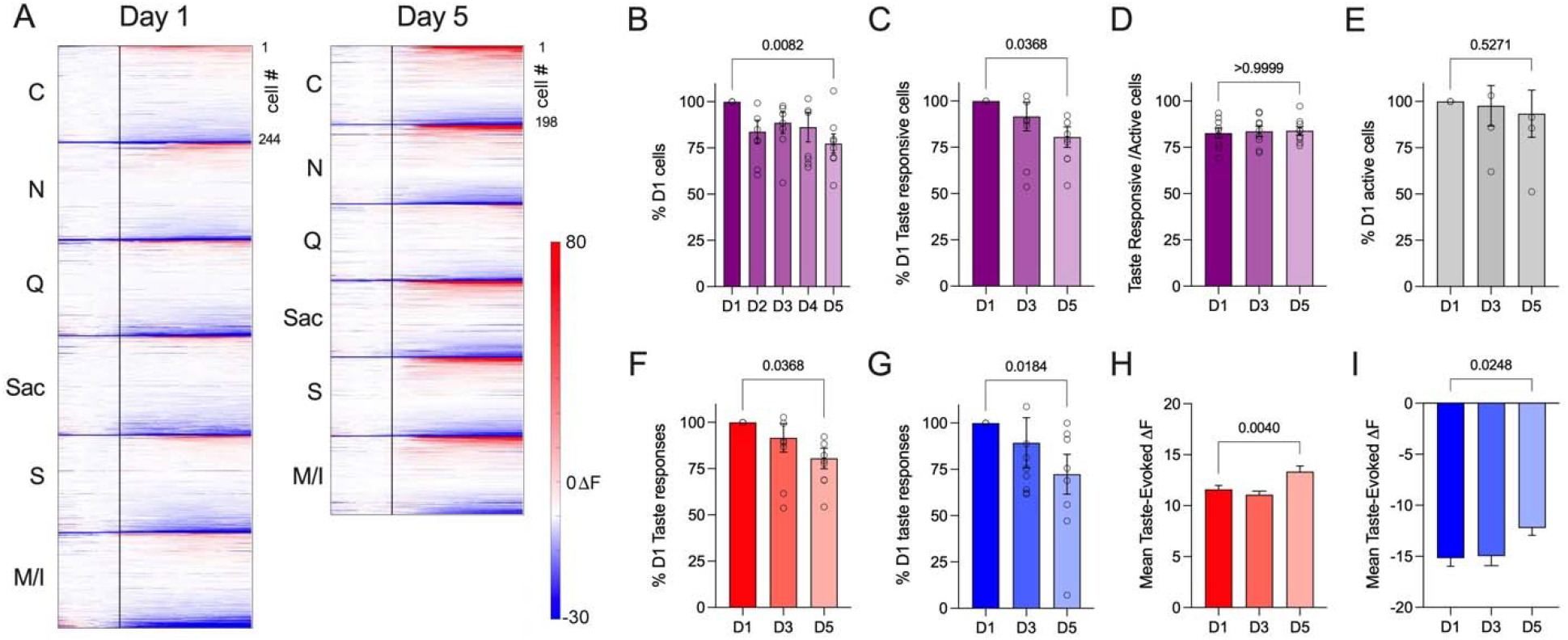
A. Individual neuron responses to the first trial of each taste taken from one mouse on day 1 and day 5. Cells are aligned by maximum to minimum DF response for each taste and day. B. Mean number of cells recorded from each mouse relative to day 1 counts. C. Percent of taste responsive neurons across days. D. The ratio of taste responsive to the total number of recorded cells per mouse across days. E. Mean number of cells recorded from each mouse on water training days relative to day 1 counts. F. Mean number of excitatory taste responses observed each day relative to day 1 counts. G. Mean number of suppressed taste responses observed each day relative to day 1 counts. H. Mean taste-evoked responses (DF) to all tastes across days in excited neurons. I. Mean taste-evoked responses (DF) to all tastes across days in suppressed neurons. Open circles in graphs depict values from individual mice. Error bars = SEM. Post hoc test p values are shown above each graph.

As both excited and suppressed taste responses were observed in the GC of each animal, we compared the number of both excited **(Figure 3F)** and suppressed taste-evoked responses **(Figure 3G)** taken from all cells observed across the five days. We found a significant day 1 to day 5 decrease in both response types (Excited: D1: 100%, D5: 80.50 ± 5.55%; Wilcoxon matched-pairs signed-rank test, p = 0.0195) (Suppressed: D1: 100%, D5: 72.37 ± 10.72%; Wilcoxon matched-pairs signed-rank test, p = 0.0391). We next compared the mean excitatory taste evoked response for all cells and tastes on each day (Excited: n=4221 total responses; Suppressed: n=683 total responses). We found a significant increase in mean taste-evoked excited responses across days (D1: 11.59 ± 0.38, D5: 13.35 ± 0.30) (Unpaired t-test t(4219)=2.678, p=0.0074) (Figure 3H). We also found a significant decrease in mean taste-evoked suppressed responses across days (D1: - 15.17 ± 0.80, D5: −12.23 ± 0.71) (Unpaired t-test t(681)=2.609, p=0.0093) **(Figure 3I).** Collectively, these data indicate that both the overall number of active cells in GC and the response characteristics of the subset of taste-responsive cells are modulated by experience.

Although we observed a decreasing number of active cells per day over our familiarization paradigm, we also identified a population of active cells that were present across taste experience. Thus, we next explored if there were any differences in response properties between cells that remained across days (i.e., “stable” cells) and ones that did not (transient cells) **(Figure 4).** On average, we found approximately 33% of the total cells per animal were stable **(Figure 4B).** Based on day 1 responses, stable cells displayed significantly stronger taste responses compared to transient cells (Mean DF: stable = 15.77 ± 0.84; transients = 9.57 ± 0.39) (Unpaired t-test t(2243)=7.612, p<0.0001) **(Figure 4C).** In contrast, no differences in mean entropy were found between the two groups in their day one responses (Unpaired t-test t(1139)=0.7939, p=0.4274) **(Figure 4D)**.

**Figure 4.**
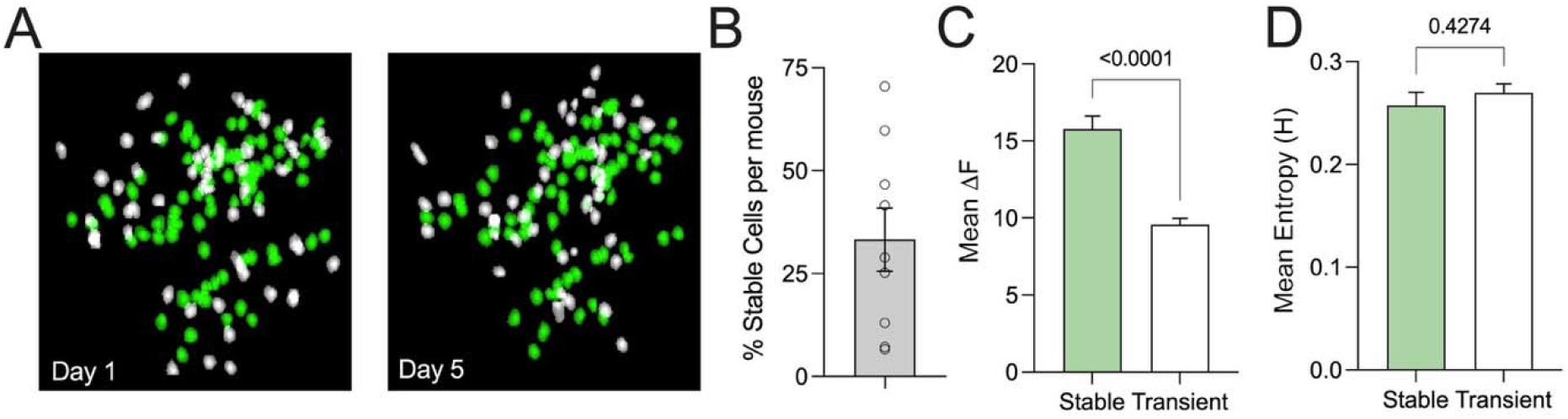
A. Cell contours were identified in the same mouse on day 1 and day 5. Stable cells are depicted in green and transient cells in white. B. Mean percent of stable cells in all mice used in the study. Open circles represent values from individual mice. C. Day 1 mean excitatory taste evoked responses (DF) in both stable and transient cells. D. Mean day 1 entropy values from stable and transient cells. Error bars = SEM. Post hoc test p values are shown above each graph.

We next chose to explore the taste coding properties of the stable population of taste-responsive cells over the course of the familiarization paradigm. We first considered how breadth of tuning in cells might change across days, using both the entropy and sparseness metrics. In all stable taste-responsive cells, entropy significantly increased across days (D1: H = 0.27 ± 0.01, D3: H = 0.30 ± 0.01, D5: H = 0.33 ± 0.01) (Kruskal-Wallis test, H(2)=24.22, p<0.0001), with post hoc Dunn’s multiple comparisons test revealing a significant difference between days 1 and 5 (p<0.0001) **(Figure 5A).** Correspondingly, we found that spareness significantly decreased across days (D1: S = 0.86 ± 0.01, D3: S = 0.84 ± 0.01, D5: S = 0.81 ± 0.01) (Kruskal-Wallis test, H(2)=21.01, p<0.0001), with post hoc Dunn’s multiple comparisons test revealing a significant difference between days 1 and 5 (p<0.0001) **(Figure 5F).** Increased entropy and decreased sparseness indicated that cells are becoming more broadly tuned over days.

**Figure 5.**
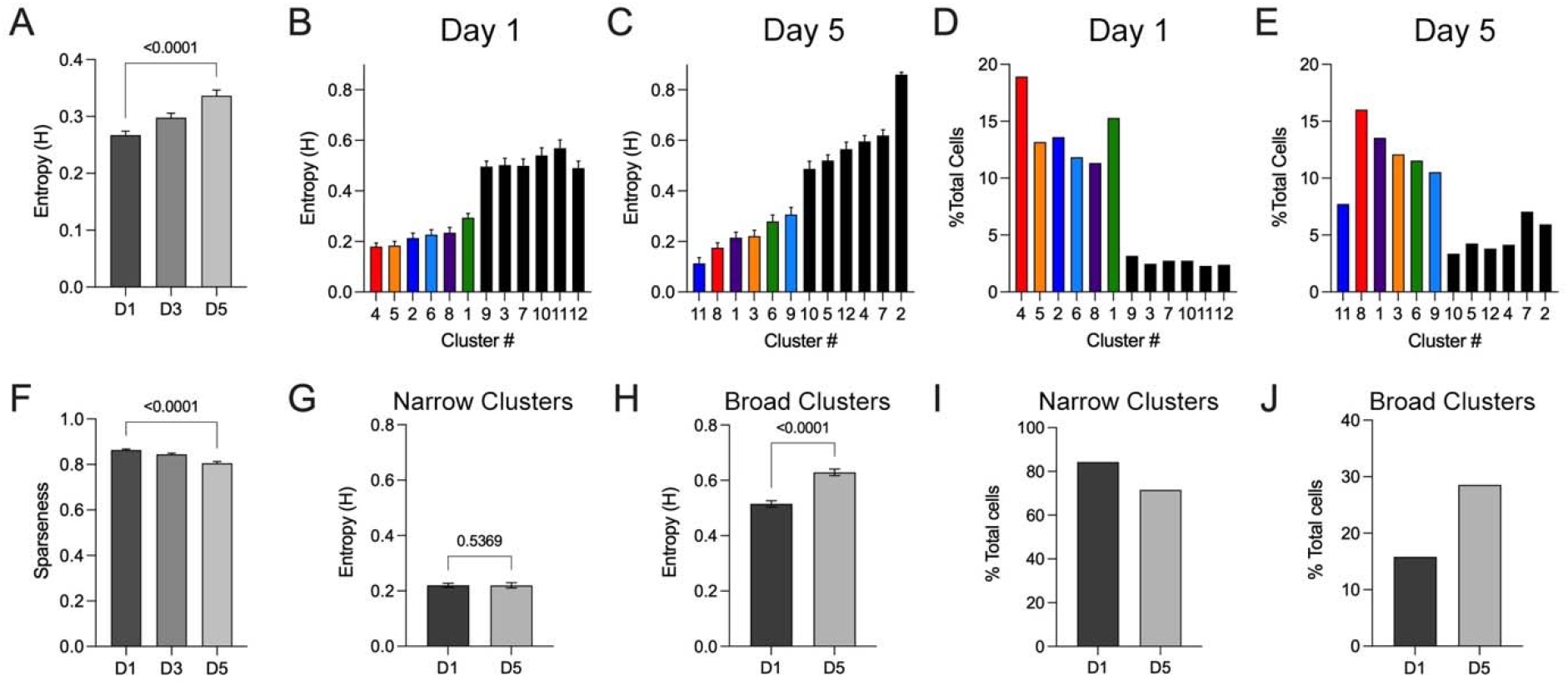
A. Mean entropy values for all stable cells across days. B, C. Mean entropy of cell clusters on day 1 (B) and day 5 (C). Narrowly tuned clusters are colored-coded by taste preference (best taste) (sucrose: red, citric acid: orange, sucrose: blue, saccharine: light blue, MPG/IMP: purple, NaCl: green). Broadly tuned clusters are depicted in black. D, E. Percentage of total taste-responsive cells in each cluster of day 1 and day 5 (E). F. Mean sparseness values for all stable cells across days. G. Mean entropy of narrow clusters on day 1 and day 5. H. Mean entropy of broad clusters on day 1 and day 5. I. Percentage of cells belonging to narrow clusters on day 1 and day 5. J. Percentage of cells belonging to broad clusters on day 1 and day 5. Error bars = SEM. Post hoc test p values are shown above each graph.

To further examine this change in tuning, we used hierarchical cluster analysis to sort cells by their taste responses and compared the clusters from day 1 **(Figure 5B)** and day 5 **(Figure 5C).** Similar to our previous 2P GC imaging study in anesthetized mice (Fletcher et al., 2017), and a more recent awake 2P imaging study (Chen et al., 2021), we found that cells sort into both narrowly tuned clusters (mean cluster entropy < 0.30) and broadly tuned clusters (mean cluster entropy > 0.40). Mean entropy values from cells in the narrow clusters were not significantly different on day 1 compared to day 5 (D1: H = 0.22 ± 0.01, D5: H = 0.22 ± 0.01) (Mann Whitney test: U=298425, p=0.5369) **(Figure 5G).** Conversely, cells in the broad clusters were found to have significantly larger entropy values on day 1 compared to day 5 (D1: H = 0.51 ± 0.01, D5: H = 0.63 ± 0.01) (Mann Whitney test: U =15281, p<0.0001) **(Figure 5H).** We also found a greater percentage of broadly tuned cells on day 5 compared to day 1 (15.75% on day 1, 28.56% on day 5). When the percentage of cells in each cluster on days 1 and 5 were compared **(Figure 5D,E),** it was evident that there was both a reduction in the percentage of cells that comprised the narrow clusters **(Figure 5I)** and an increase in the percentage cells that comprise the broad clusters **(Figure 5J)**.

As cells were found to increase their breadth of tuning across days, we next tested whether this increase might lead to changes in the similarity of population representations of each taste. We pooled all excited taste responses for all mice on days 1, 3, and 5 and performed pair-wise correlations between responses for each taste. Strikingly, the mean taste-taste response correlations increased over days between all appetitive tastes **(Figure 6A).** To further test this, we compared the mean of all response correlations between all stimuli (except quinine) on each day and found that mean correlations significantly increased across days (D1: 0.12 ± 0.02; D3: 0.23 ± 0.06; D5: 0.75 ± 0.02) (Friedman’s test, χ2(3) = 20.13, p<0.0001) Post-hoc analysis revealed a significant increase between days 1 and 5 (Dunn’s multiple comparison test: p<0.0001) and between days 3 and 5 (Dunn’s multiple comparison test: p=0.0016) suggesting that response similarity occurs after at least three days of familiarity trials **(Figure 6B).** While mean licking to all stimuli (except quinine) does significantly increase over days (D1: 28.91 ± 1.81; D3: 38.81 ± 1.01; D5: 37.28 ± 1.27) (ANOVA, F(2,106) = 26.77, p<0.0001) **(Figure 6D),** this effect cannot fully account for the increased response correlations as increases in licking are seen between days 1 and 3 (Bonferonni’s multiple comparison test: p<0.0001), but not between days 3 and 5 (Bonferonni’s multiple comparison test: p<0.8819). Interestingly, while response correlations between appetitive stimuli increased, response correlations to between appetitive stimuli and quinine remained stable over the five days (D1: 0.06 ± 0.01; D3: 0.11 ± 0.01; D5: 0.13 ± 0.05) (Friedman’s test, χ2(3) = 3.00, p<0.2522) **(Figure 6C).** Further, no changes in licking on quinine trials was observed across days (D1: 5.57 ± 1.13; D3: 4.86 ± 2.92; D5: 3.43 ± 1.95) (ANOVA, F(2,12) = 26.77, p<0.0001) **(Figure 6E)**.

**Figure 6.**
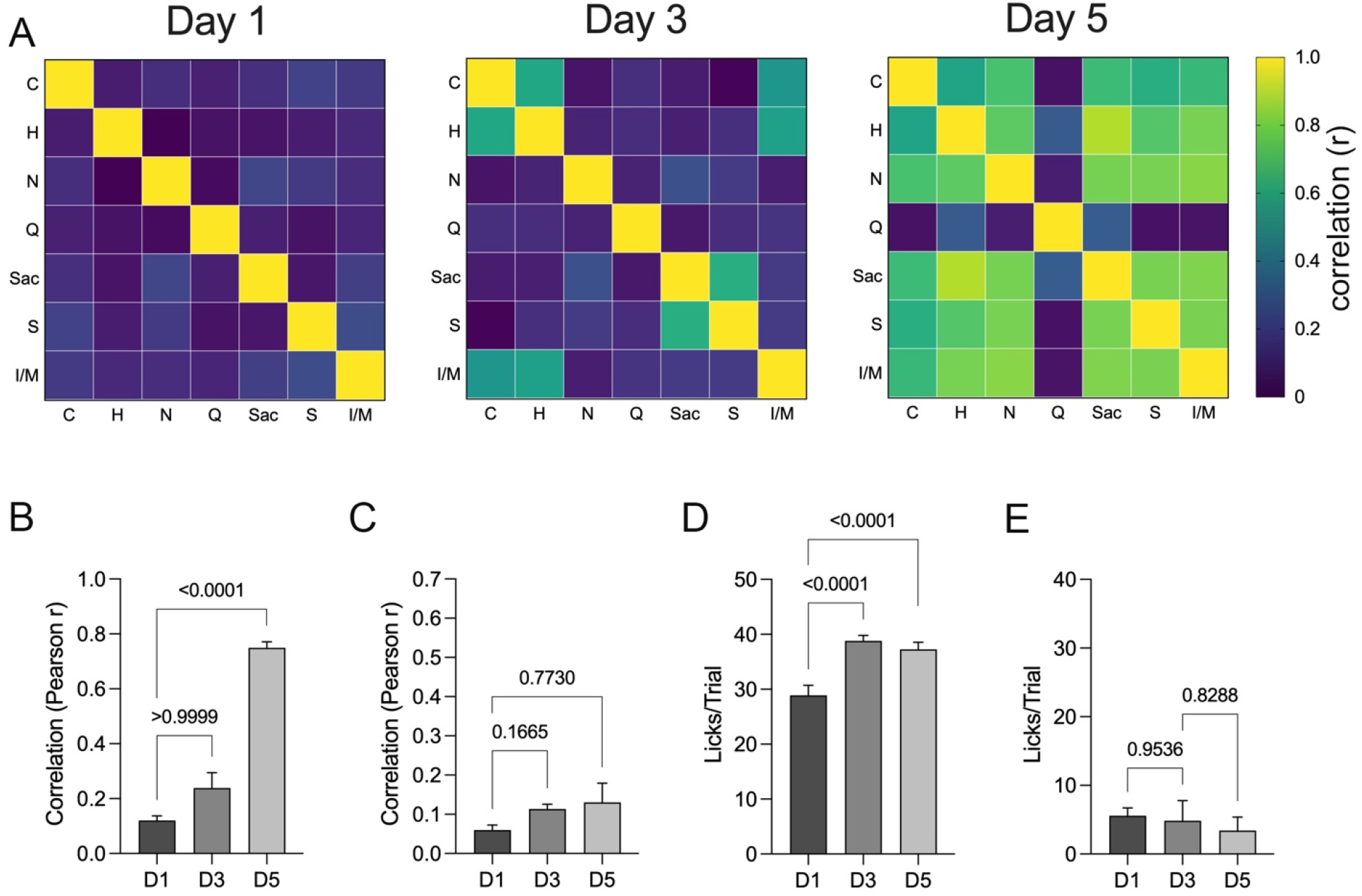
A. Correlation matrices for pooled responses to all tastes on days 1, 3, and 5. B. Mean response correlations between all stimuli (except quinine) on each day. C. Mean response correlations between each appetitive stimuli and quinine on each day. D. Mean licks per trial to all stimuli except quinine on each day. E. Mean licks per trial quinine on each day. Error bars = SEM. Post hoc test p values are shown above each graph.

## Discussion

### Imaging taste responses in GC neurons freely moving mice

In this study, we investigated how taste familiarity impacts GC neuron activity and taste coding in mice. Head-mounted miniaturized microscopes were used in conjunction with an implanted GRIN lens to visualize calcium activity within individual GC neurons across multiple days of taste exposure in freely-moving mice, sampling taste solutions in a lickometer. This design is different from other recent GC imaging or physiological studies of taste coding with awake, behaving mice, where mice either received the stimuli via intraoral cannula (Levitan et al., 2019), or were head-fixed and trained (over multiple sessions) to lick stimuli from a spout upon association with an auditory cue (Bouaichi and Vincis, 2020; Chen et al., 2021). We chose our experimental design to more closely approximate how taste/ingestive behavior in mice is typically measured using brief-access trials in standard lickometers (Boughter et al., 2005; Schier et al., 2019). The lack of taste training also allowed us to examine responses to stimuli as they progress from novel to familiar across test sessions. On the other hand, our design introduces a number of other factors inherent to the lickometer that could impact the taste response, such as trial-to-trial variability in the latency to approach the spout, variability in the amount of fluid consumed in each trial, and behaviors that occur in between trials, such as grooming and exploration of the test chamber. Crucially, a comparison of lick responses of miniscope mice vs. non-surgical controls revealed that the miniscopes themselves did not appear to impact brief-access taste behavior.

Despite these differences in behavioral design, the basic response properties of GC neurons to taste stimuli were generally consistent with the other recent studies in mice. Across all animals and days, a majority of GC neurons responded to at least one stimulus, with most (>80%) of those responses being excitation. Furthermore, we found similar distributions of taste responses across all mice and no overrepresentation of any one taste in any mouse, lending further support to further the idea of non-topographic, distributed taste identity coding within GC (Fletcher et al., 2017; Lavi et al., 2018; Levitan et al., 2019; Bouaichi and Vincis, 2020; Chen et al., 2021). Although clusters of both narrowly and broadly tuned cells were apparent in the dataset, the majority of GC neurons were characterized as narrowly tuned. This finding was similar to our previous anesthetized imaging study (Fletcher et al., 2017) and a more recent awake imaging study (Chen et al., 2021), but different from recent awake multiunit recording studies in mice, where a greater proportion of individual neurons possessed broad response profiles (Levitan et al., 2019; Bouaichi and Vincis, 2020). It is unclear whether this variation is related to the recording method (imaging vs. physiology), or perhaps even to the behavioral paradigm employed.

### The number of responsive neurons in GC is modulated by familiarization

Over the course of 5 days of familiarization, we found a significant decrease in the number of active, identifiable GC cells. This decrease was not primarily driven by a reduction in cells characterized as taste-responsive, as the overall proportion of active cells to taste-responsive cells remains similar across days. Furthermore, the effect was not likely due to cells merely dropping out of the imaging field of view with time, as active cell number remained stable across an identical number of pre-taste days in which mice only drank water. Prior immunocytochemical studies reported reduced numbers of activated neurons (i.e., c-Fos or pERK expression) in animals consuming familiar tastes versus novel tastes (Lin et al., 2012; Bamji-Stocke et al., 2018; Kayyal et al., 2021). In each of these studies, the familiar condition was achieved by allowing animals 6 or more days of exposure prior to perfusion. Similarly, in our experiment, the number of responsive cells diminished across days, with a significant Day 1 versus Day 5 difference, suggesting more cells are engaged in GC when tastes are novel. One possibility worth considering is that the effect found here is related to taste neophobia, a phenomenon where rodents are hesitant to consume novel taste stimuli (even those classified as appetitive), but increase consumption upon subsequent exposures as the novel stimulus proves safe to ingest (Domjan, 1976; Reilly, 2018). This jump in consumption is typically most conspicuous from day 1 to day 2 or 3, after which behavior asymptotes (Domjan and Gillan, 1976; Gutierrez et al., 2003; Arthurs et al., 2018; Shinohara and Yasoshima, 2019). Compared to our familiarization paradigm, taste neophobia experiments generally use a single novel tastant, as expression of neophobia to any one stimulus is diminished with pre-exposure to, or experience with, multiple stimuli (Domjan, 1977; Braveman, 1978). Even so, it was evident that lick counts to most stimuli increased across days in our miniscope mice, especially from day 1 to day 2. Licking behavior was fairly stable from day 2-5, whereas the number of taste-responsive cells declined across the same time frame, suggesting that attenuation of neophobia may not fully account for the neuronal effect.

Another possibility for the reduction in responsive cell number is that this effect reflects a process of sensory habituation, whereby repeated exposure to stimuli without other consequences results in diminished behavioral response with concomitant attenuated neural response in the CNS (Weinberger, 1995). For example, neuronal responses in the olfactory bulb have been shown to habituate following repeated daily odor exposure (Fletcher, 2012; Kato et al., 2012; Ross and Fletcher, 2018). Similarly, Kato et al. (Kato et al., 2015) found that daily experience with a simple auditory tone caused progression habituation over days of the cortical representation of the tone, including a decreased number of layer 2/3 excited cells (with a concomitant increase in inhibited cells). This withdrawal of excitation is similar to our results in terms of timing, although we also found a decrease in inhibited cells. There has been a relative lack of similar studies at the neuronal level in the taste CNS, although it has been shown that mice will adapt behaviorally to taste stimuli, e.g., becoming less sensitive to bitter-tasting stimuli after several weeks of experience (Mura et al., 2018). Bahar et al. (Bahar et al., 2004) recorded multiunit activity in the GC in rats actively sampling a single taste stimulus and water over three consecutive days. They reported that the average response to the stimulus actually increased as the taste became familiar, although this change was only apparent in the late phase of the response. In contrast, Flores et al. (Flores et al., 2022) found no change in overall GC responsiveness with three days of familiarization training with two stimuli and water, but did find changes in how these stimuli were encoded (see below). These studies necessarily focused on the responses of only taste-responsive cells. Although the overall number of taste-responsive cells in our study dropped across days, we also found that the mean taste-evoked excited response in those cells increased.

### Familiarity alters GC taste coding

Previous studies using awake recordings in GC have amply demonstrated how associative learning, or even changes in attentional state, shapes taste coding both on the single-neuron and population level (Fontanini and Katz, 2006, 2008; Vincis and Fontanini, 2016). For example, training rats to associate a non-taste cue with a taste stimulus leads to a more rapid temporal encoding of stimuli in GC neurons (Samuelsen et al., 2012; Mazzucato et al., 2019). This includes a decrease in entropy values (i.e., more narrow tuning) in neurons when stimuli were expected rather than unexpected. In our data, we examined potential changes in taste coding in the population of stably responding cells during taste familiarization. On average, stable cells actually became more broadly tuned across days, both in terms of entropy and sparseness. This increase was not seen in all individual cells; rather, the percentage of neurons classified (via cluster analysis) as broad responders increased from Day 1 to Day 5. This difference from the Samuelsen et al. (Samuelsen et al., 2012) study may be related to the lack of associative learning used in our study. Further analysis of our data indicated that most tastes became progressively more correlated in terms of neural response across days. The exception to this trend was quinine, which remained uncorrelated to the other tastes, maintaining a distinct neural pattern. Unsurprisingly, quinine was also the outlier in licking behavior; it was strongly avoided throughout the behavioral paradigm, whereas even citric acid was licked at a rate similar to water, sucrose, or saccharin, especially by days 4-5. The increase in correlation among stimuli excepting quinine was likely not simply due to neural activity reflecting the number of licks or amount of fluid in the oral cavity. First, although mice only lick quinine a few times in a trial compared to the other stimuli (even on day 1), a similar percentage of cells responded to all six stimuli, and the mean quinine-evoked excited response was the highest among stimuli. Second, although lick counts to all appetitive stimuli were similar on days 3 and 5, the interstimulus correlation increased dramatically as compared on those two days. On a side note, the increase in licking to the sour stimulus citric acid (0.02 M) over days was somewhat surprising. It is not as inherently aversive to mice as 0.01 M quinine (Fletcher et al., 2017), and we are unaware of studies examining licking to citric acid over time.

These results indicate that at least in the population of stable taste-responsive GC cells, neural responses evolve from a pattern of dissimilarity to all novel taste stimuli, to one where stimuli evoking similar behavioral responses elicit similar neural patterns. At first glance, these results seem at odds with a recent physiological study in GC in rats, where 3 days of experience with NaCl, citric acid, and water led to an increase in the distinctness of the neural patterns evoked by these stimuli (Flores et al., 2022). However, stimulus delivery was via IO cannulas, so the specific behaviors evoked by the pair of tastants are inferred – it is possible that the citric acid remains aversive over days to rats, unlike the change seen in mice. Furthermore, in both this and the earlier study, changes over days were found in the late phase response, i.e.,> 1 s after drinking onset (Bahar et al., 2004; Flores et al., 2022); by this time point, GC neurons are encoding a palatability-related response (Katz et al., 2001; Levitan et al., 2019). It is possible, then, that the increased correlations among avidly licked stimuli found here (and the lack of correlation with quinine) represent a palatability-related population response that increases in strength across days. Although imaging generally lacks the temporal precision needed to define response periods on a millisecond scale as in the physiology work, previous GC imaging work noted that palatability information can be discerned in the population response (Fletcher et al., 2017). Follow-up experiments might test this idea more explicitly, i.e. test correlation over time with a panel of tastes including additional aversive taste stimuli, or testing the effect of making a palatable stimulus unpalatable through associative learning such as CTA (following familiarization). The latter idea has actually been examined in previous studies, though often with different approaches. For example, following CTA, saccharin evoked a quinine-like pattern of activity on the cortical surface as measured by intrinsic signal imaging (Accolla and Carleton, 2008). Using 2P imaging of neurons in GC (including those that project to the amygdala), Lavi et al. (Lavi et al., 2018) showed that CTA shifted the representation of a sweet conditioned stimulus (also saccharin) from uncorrelated to highly correlated with quinine.

While many studies of taste physiology rely on repeated exposures to taste stimuli, little work has focused on how the process of taste experience itself may alter cortical taste responses. Overall, the results of our experiments indicate at least two neuronal processes occurring during familiarization with a panel of tastants. First, GC becomes less “noisy” with repeated experience in terms of a drop in active cells across days. Some of these cells may have functions other than taste, such as attentional state, but taste-responsive neurons are also reduced in number with experience. These findings are best explained in terms of sensory habituation, described in other sensory cortical areas (Weinberger, 1995; Quairiaux et al., 2007; Kato et al., 2015). In the subpopulation of stable GC neurons that are present on each test, however, the population response to tastes evolves to reflect stimulus similarities in terms of behavior. While few previous studies have examined neuronal changes with taste familiarization, multiple behavioral studies have documented this process of familiarization learning has important consequences for subsequently learned behaviors. For example, prior exposure to a particular taste stimulus reduces the ability of an animal to form a CTA to that (conditioned) stimulus, a process known as latent inhibition (Garcia et al., 1974; Clark and Bernstein, 2009; Lubow, 2009). Conversely, familiarization with several novel tastants can actually enhance CTA to a different, novel conditioned stimulus (Flores et al., 2016; Flores et al., 2018), as well as impact the response to a novel unconditioned stimulus (Flores et al., 2022). This enhancement has been shown to be dependent on the GC (Flores et al., 2018). Future experiments might focus on the effect of taste familiarization on specific subpopulations of GC cells, either those expressing distinct molecular markers or with specific projection patterns (Kato et al., 2015).

## Author Contributions

Conceptualization: M.L.F., S.M.S., and J.D.B.; Methodology, M.L.F., S.M.S., and J.D.B.; Data Collection: S.M.S.; Formal analysis, S.M.S. and M.L.F. Original Draft, S.M.S., M.L.F. and J.D.B.

## Acknowledgements

The Optogenetics and Neural Engineering (ONE) Core at the University of Colorado School of Medicine provided engineering support for this research. The ONE Core is part of the NeuroTechnology Center, funded in part by the School of Medicine and by the NINDS of the National Institutes of Health under award number P30NS048154. This study was supported by NIH grant NIDCD DC016833 to M.L.F. and J.D.B.

## Notes

### Competing Interest Statement

The authors have declared no competing interest.

