## Supplemental Figures for "The impact of familiarity on cortical taste coding"

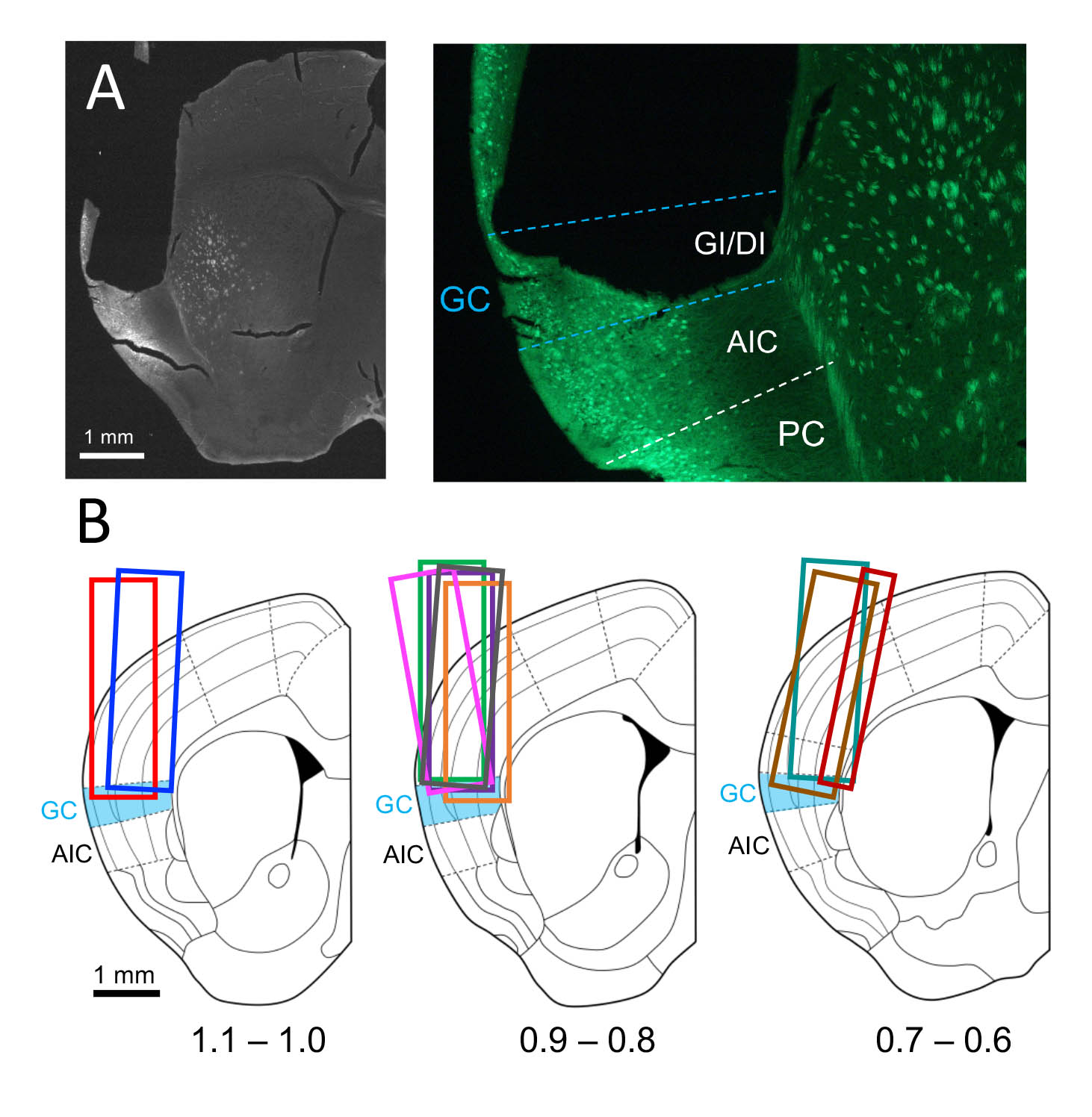

**Supplemental Figure 1.** Histological placement of GRIN lenses. A. Representative lens placement, shown at lower and higher power. The outline of the GRIN lens can be seen in the left cortical hemisphere, just above GC. GCaMP6s expression is expressed in this region. B. Schematic showing the location and A-P level of GRIN lenses in all 9 mice. Each color designates a different animal. All lenses were 1 mm in diameter except for one 0.5 mm lens, shown in dark red. Abbreviations: GI/DI = granular/dysgranular cortex, which corresponds approximately to gustatory cortex (GC). AIC = agranular insular cortex, PC = piriform cortex.

**Supplemental Figure 2.** Comparison of mean lick ratio (number of licks to the tastant/mean licks of water trials) per trial for each taste across five days of behavior in miniscope mice (n=9)(magenta) and non-surgical control mice (n=12) (black). No statistical differences were found across groups for any of the tastants (2-way ANOVA). Error bars = SEM.

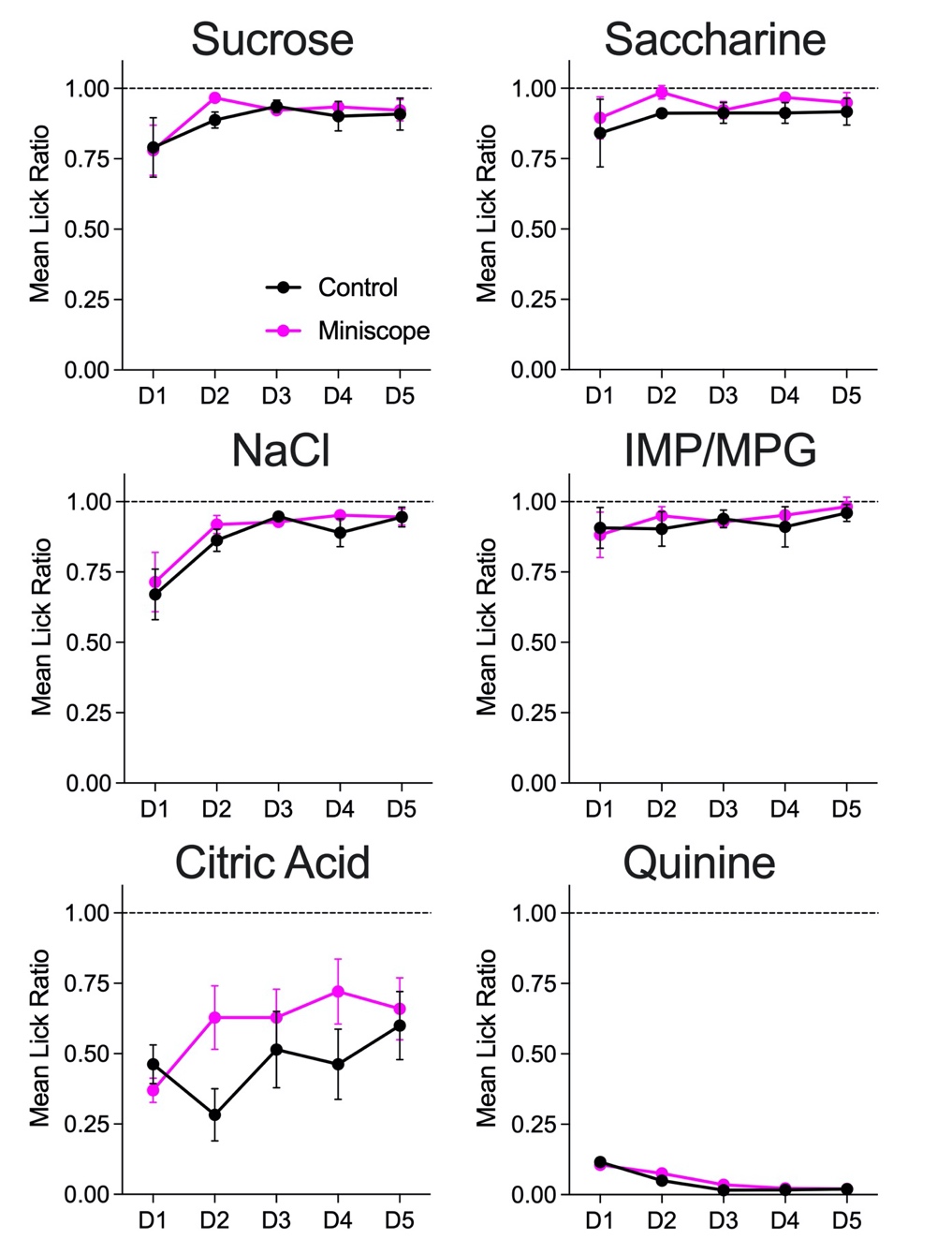
